# The Boundary-Expressed *EPIDERMAL PATTERNING FACTOR-LIKE2* Gene Encoding a Signaling Peptide Promotes Cotyledon Growth during *Arabidopsis thaliana* Embryogenesis

**DOI:** 10.1101/2021.04.27.441542

**Authors:** Rina Fujihara, Naoyuki Uchida, Toshiaki Tameshige, Nozomi Kawamoto, Yugo Hotokezaka, Takumi Higaki, Rüdiger Simon, Keiko U Torii, Masao Tasaka, Mitsuhiro Aida

## Abstract

The shoot organ boundaries have important roles in plant growth and morphogenesis. It has been reported that a gene encoding a cysteine-rich secreted peptide of the EPIDERMAL PATTERNING FACTOR-LIKE (EPFL) family, *EPFL2*, is expressed in the boundary domain between the two cotyledon primordia of *Arabidopsis thaliana* embryo. However, its developmental functions remain unknown. This study aimed to analyze the role of *EPFL2* during embryogenesis. We found that cotyledon growth was reduced in its loss-of-function mutants, and this phenotype was associated with the reduction of auxin response peaks at the tips of the primordia. The reduced cotyledon size of the mutant embryo recovered in germinating seedlings, indicating the presence of a factor that acted redundantly with *EPFL2* to promote cotyledon growth in late embryogenesis. Our analysis indicates that the boundary domain between the cotyledon primordia acts as a signaling center that organizes auxin response peaks and promotes cotyledon growth.

## Introduction

The growth and morphogenesis of plant organs are manifested by the interactions between each organ and surrounding regions. An important class of developmental domains that affects the shoot organ development is the boundary domain, which is generated between the adjacent shoot organ primordia or between the shoot meristem and organ primordium (Aida and Tasaka 2006). The boundary domain is characterized by the expression of specific regulatory genes and differential activities of plant hormones. It also plays pivotal roles in shoot meristem activity regulation, adjacent shoot organ separation, and new shoot meristem formation (Hepworth and Pautot 2015; Žádníková and Simon 2014).

The interactions between the different developmental domains are partly mediated by signaling molecules, such as hormones and small peptides. Among these molecules, the EPIDERMAL PATTERNING FACTOR-LIKE (EPFL) family of secreted cysteine-rich proteins is involved in various developmental pathways, including epidermal cell patterning, inflorescence architecture, and lateral shoot organ patterning (Tameshige et al. 2017; Torii 2012). Several studies have shown that one of the genes encoding an EPFL family member, *EPFL2*, is specifically expressed in the boundary domains of various shoot organs and regulates shoot meristem size, leaf and ovule positioning, and leaf margin morphogenesis (Kawamoto et al. 2020; Kosentka et al. 2019; Tameshige et al. 2016). The boundary-specific expression of this gene has also been reported during embryogenesis (Kosentka et al. 2019), in which a series of patterning and growth events occur to establish the basic body plan (Palovaara et al. 2016).

Although the functions of *EPFL2* in the boundary domain have been investigated in various developmental contexts, the role of this gene in embryogenesis is unknown. This study aimed to investigate the function of *EPFL2* during embryogenesis. The expression analysis was extended to earlier embryonic stages relative to the previous report by Kosentka et al. (2019), and we found that this gene was expressed in the embryo apex before the initiation of cotyledon primordia. The analysis of loss-of-function mutants of *EPFL2* showed that the size of the cotyledon primordia was significantly reduced, and the activity of an auxin response marker, *DR5*, was also significantly decreased. The results suggest that the boundary domain in the embryo apex plays an important role in promoting the growth of adjacent cotyledons.

## Materials and Methods

### Plant materials and growth conditions

The *A. thaliana* accessions L*er* and Col were used as the wild-type strains. The *EPFL2pro::GUS* reporter (Col background), transposon insertion allele *epfl2-1* (L*er* background), and CIRSPR/Cas9-induced alleles *epfl2-2* and *epfl2-3* (Col background) were described previously (Kawamoto et al. 2020; Tameshige et al. 2016). For the analysis of *DR5rev::GFP* (Friml et al. 2003), the *epfl2-1* allele was backcrossed to Col seven times and then crossed with the reporter line of the Col background (Tameshige et al. 2016). The plants were grown as described previously (Takeda et al. 2011).

### Microscopy

GUS staining was done as previously described (Aida et al. 2020). For the visualization of the embryo morphology, the ovules were cleared as described previously (Aida et al. 1999), and their stages were determined according to Jürgens and Mayer (1994). To measure the length and cell number of the cotyledons in germinating seedlings, the seeds were first imbibed on wet filter paper for three days at 4 °C in the dark; then, they were incubated in a growth chamber at 23 °C under constant white light for 24 h. After removing the seed coat, the cotyledons were excised and directly mounted in a clearing solution (8 g of chloral hydrate, 1 ml of glycerol, and 2 ml of water). The measurements were carried out in the palisade layer. For the analysis of *DR5rev::GFP*, the embryos were cleared as described previously (Imoto et al. 2021), and the confocal images were taken using LSM 5 Live (Zeiss [Oberkochen, Germany]). Because the signal intensities were generally much higher in the root than in the cotyledons (∼5.7 fold), a pair of images with different fixed values of Main Gain parameter (50 and 25 for cotyledons and root, respectively) was independently collected for each embryo to avoid the saturation of the signals of interest. To quantify the GFP signals, the average background value in a small area within the embryo was subtracted from the maximum signal value in the protoderm of the cotyledon tips or that in the outermost columella root cap cells. The image and statistical analyses were performed using Fiji (Schindelin et al. 2012) and R (version 3.6.1; The R Foundation for Statistical Computing Platform), respectively.

## Results

### Expression of *EPFL2* during embryogenesis

We first examined the expression patterns of the *EPFL2* gene in the embryo using a GUS reporter. The GUS activity was first detected in a small area of the apical region at the mid-globular stage (Figure 1A). The embryos with an oblique view indicate that the expression was initiated as a pair of asymmetric spots (Figure 1A, inset). At the heart stage, the expression was detected in the boundary domain between the two cotyledon primordia (Figure 1B, C), and this expression pattern continued in the later stages (Figure 1D). Within the boundary domain, the GUS activity was much stronger in the periphery than in the center (Figure 1C). These results indicate that the expression of *EPFL2* starts before the cotyledon initiation and continues in the boundary domain in the later stages of embryogenesis.

**Figure 1.**
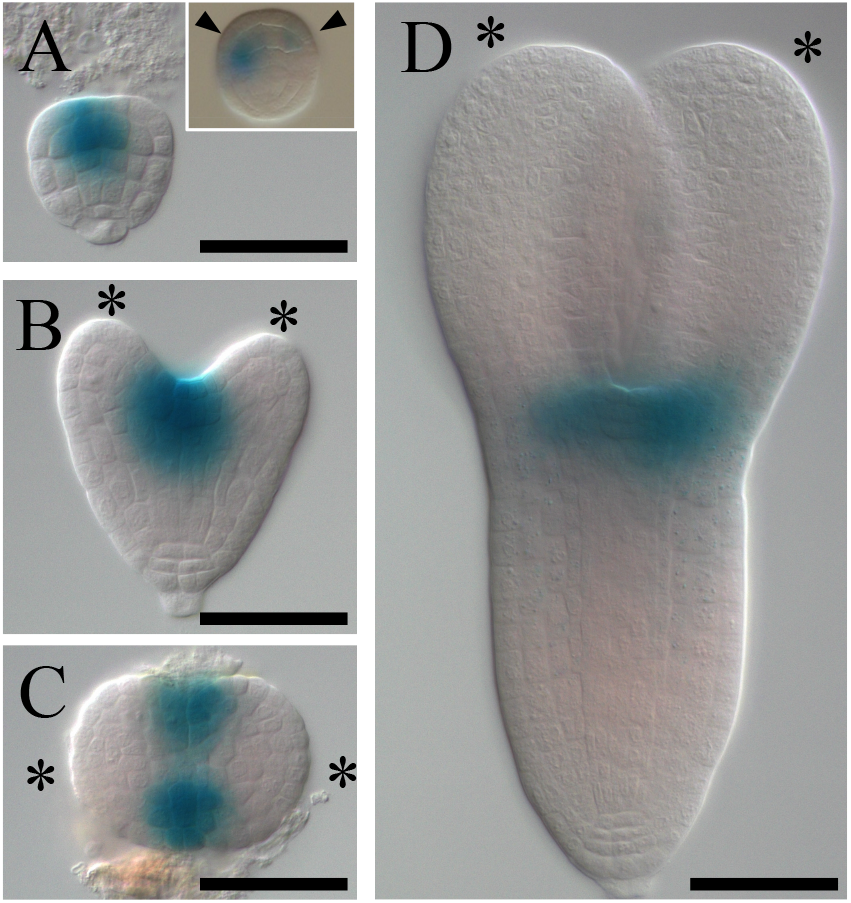
Expression patterns of *EPFL2pro::GUS*. Wild-type embryos at the mid-globular (A), mid-heart (B and C), and mid-torpedo (D) stages. Frontal (A, B, and D) and top (C) views. The inset in (A) shows an embryo with two asymmetric spots of GUS activity (arrowheads) on oblique view. The asterisks indicate the cotyledon primordia. Scale bars = 50 μm.

### *EPFL2* is required for cotyledon growth during embryogenesis

To investigate the function of *EPFL2*, we examined the effect of its loss-of-function mutations on embryo development. The null allele *epfl2-1* (Tameshige et al. 2016), which was generated in the L*er* background, did not display obvious morphological defects up to the globular stage (Figure 2A, B). However, after the heart stage, the cotyledon primordia were shorter compared to those of the wild-type L*er* (Figure 2C, D). We quantified this phenotype and found that the height of the cotyledon primordia relative to that of the rest of the embryo (referred to as cotyledon height and axis height, respectively; Figure 2E) was smaller in *epfl2-1* than in L*er* (Figure 2F). The reduction of the cotyledon height was the most prominent when the axis height was 51–100 μm (47.7 % reduction; Figure 2G), and it became less as the axis height increased (33.7 % in 101–150 μm and 32.2 % in 151–200 μm; Figure 2G).

**Figure 2.**
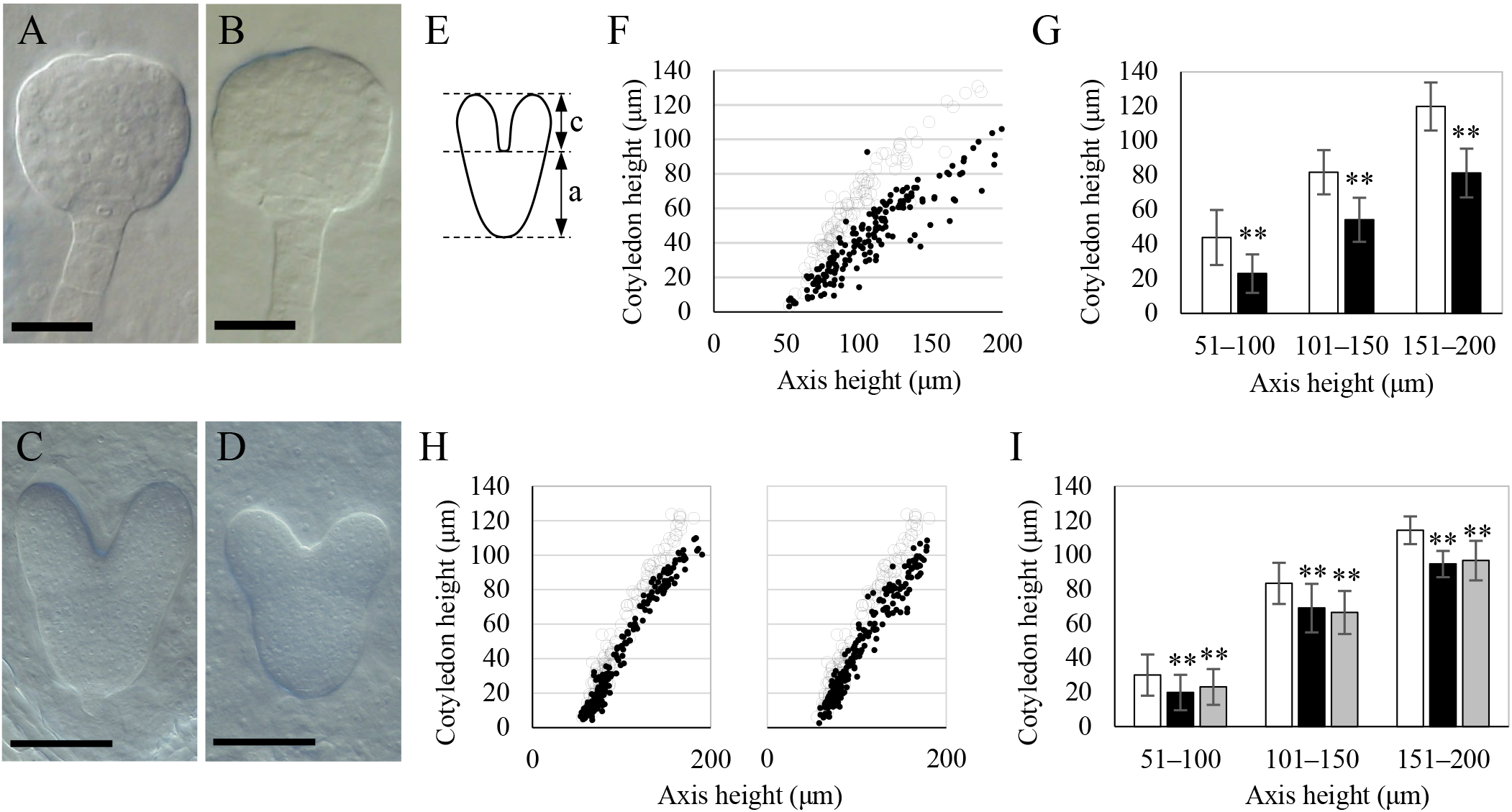
Embryonic phenotype of *epfl2* mutants. (A to D) Cleared embryos at the mid-globular stage of wild-type L*er* (A) and *epfl2-1* (B) and those at the late-heart stage of L*er* (C) and *epfl2-1* (D). (E) Quantification of the cotyledon size. The height of the cotyledon (c) and axis (a) was measured. The axis refers to the region that spans the shoot apex and the root tip. (F) Relationship between the axis and cotyledon height. The open circles and black dots represent the individual embryos of L*er* and *epfl2-1*, respectively. (G) Cotyledon height of embryos with three classes of different axis height. The open and closed bars represent L*er* and *epfl2-1*, respectively. (H) Relationship between the axis and cotyledon height of wild-type Col vs *epfl2-2* (left panel) and Col vs *epfl2-3* (right panel). The open circles and black dots represent the individual embryos of Col and mutants, respectively. Each mutant data set is compared to the same data set of Col. (I) Cotyledon height of embryos with three classes of different axis height. The open, closed, and gray bars represent Col, *epfl2-2*, and *epfl2-3*, respectively. Scale bars = 20 μm in A and B and 50 μm in C and D. The three classes with different axis height roughly correspond to heart (51–100 μm), early-torpedo (101– 150 μm), and mid-torpedo (151–200 μm) stages according to Jürgens and Mayer (1994). The error bars represent the standard deviation. The double asterisks indicate the significant differences between each mutant and wild-type control (p < 0.01; Welch’s t-test for G and Dunnett’s test for I). The sample sizes for each measurement are described in Supplementary File 1.

We also analyzed *epfl2-2* and *epfl2-3*, which are CRIPR/Cas9-induced alleles in the Col background, and they both showed similar reduction in cotyledon height (Figure 2H, I), although their phenotypes were milder compared to that of the L*er* allele *epfl2-1* (e.g., 33.9 % and 23.0 % reduction in *epfl2-2* and *epfl2-3*, respectively, for embryos with 51– 100 μm axis height range). The deduced amino acid sequence produced from *epfl2-2* completely lacks a mature peptide (Kawamoto et al. 2020), indicating that this allele is null, similar to *epfl2-1*. Therefore, the observed difference in the phenotypic severity of *epfl2-1* and *epfl2-2* is likely due to their difference in genetic background rather than that in allele strength. Besides the size reduction in cotyledon primordia, no obvious abnormality was found in any of the *epfl2* mutant alleles. Our analysis shows that *EPFL2* is required to promote cotyledon growth during embryogenesis.

### The reduction in cotyledon size of *epfl2* mutants was rescued in germinating seedlings

We also examined the early seedling phenotypes of *epfl2* mutants. The shape and size of the cotyledons in *epfl2* mutants were indistinguishable from those of the wild type (Figure 3A, B). The formation of leaves also occurred normally, except that the leaf margins lacked serrations, as reported previously (Tameshige et al. 2016). The quantification of the length and cell number along the proximo-distal and lateral axes of the cotyledons showed no significant reduction in these values in germinating, one-day-old seedlings of the two Col alleles (Figure 2C, D). In *epfl2-3*, the size and cell number along the proximo-distal axis were significantly greater than those of Col. Although the reason for this allele-specific effect is unclear, our results show that neither of the *epfl2* mutations caused the reduction in cotyledon size or cell number, indicating that the reduced cotyledon growth in mutants during embryogenesis was rescued in germinating seedlings.

**Figure 3.**
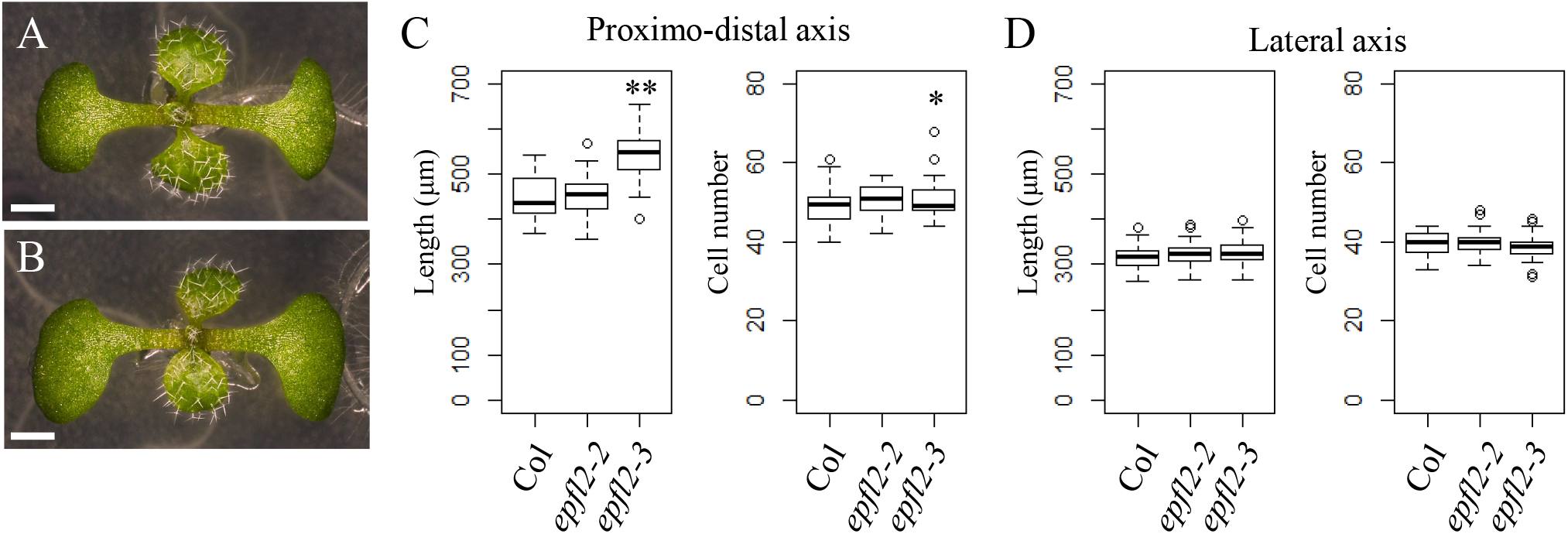
Seedling phenotype of *epfl2* mutants. (A and B) Top views of seven-day-old wild-type Col and *epfl2-2* mutant seedlings. (C) Length (left) and cell number (right) of cotyledons along the proximo-distal axis in one-day-old wild-type Col and *epfl2* mutant seedlings. (D) Length (left) and cell number (right) of cotyledons along the lateral axis in one-day-old Col and *epfl2* mutants. The single and double asterisks indicate the significant differences between each mutant and wild-type control (p < 0.05 and p < 0.01, respectively; Dunnett’s test for C and D). Scale bars = 1 mm. The sample sizes for each measurement are described in Supplementary File 1.

### *EPFL2* is required to establish auxin response peaks at the cotyledon tips

It has been established that the formation of auxin response peaks (also called auxin maxima) is essential for proper cotyledon development (Benková et al. 2003). Therefore, we analyzed the patterns of the auxin response reporter, *DR5rev::GFP*, in *epfl2* embryos (see Materials and Methods). In the wild type, the GFP signals were detected as a pair of apical spots and single basal spot, each of which corresponds to the *DR5* activity at the two cotyledon tips and root tip, respectively (Figure 4A). In the wild type, most embryos possessed two recognizable apical signals at the heart stage. However, within a single embryo, the signal intensity was often different in each of the cotyledon tips (arrowheads in Figure 4A). In contrast, the *epfl2* mutant embryos often lack recognizable apical signals in one or both of the cotyledon tips (Figure 4B, C). The intensity of the apical signals was significantly lower in *epfl2* than in the wild type (60.9 % reduction; Figure 4D). In contrast to the reduction of the apical signals, the *epfl2* mutant embryos did not show significant changes in the pattern or intensity of the GFP signals in the root pole (Figure 4A, B, E). These results demonstrate that *EPFL2* is required to establish auxin response peaks in the apical embryo, and are consistent with the specific role for this gene in cotyledon development during embryogenesis.

**Figure 4.**
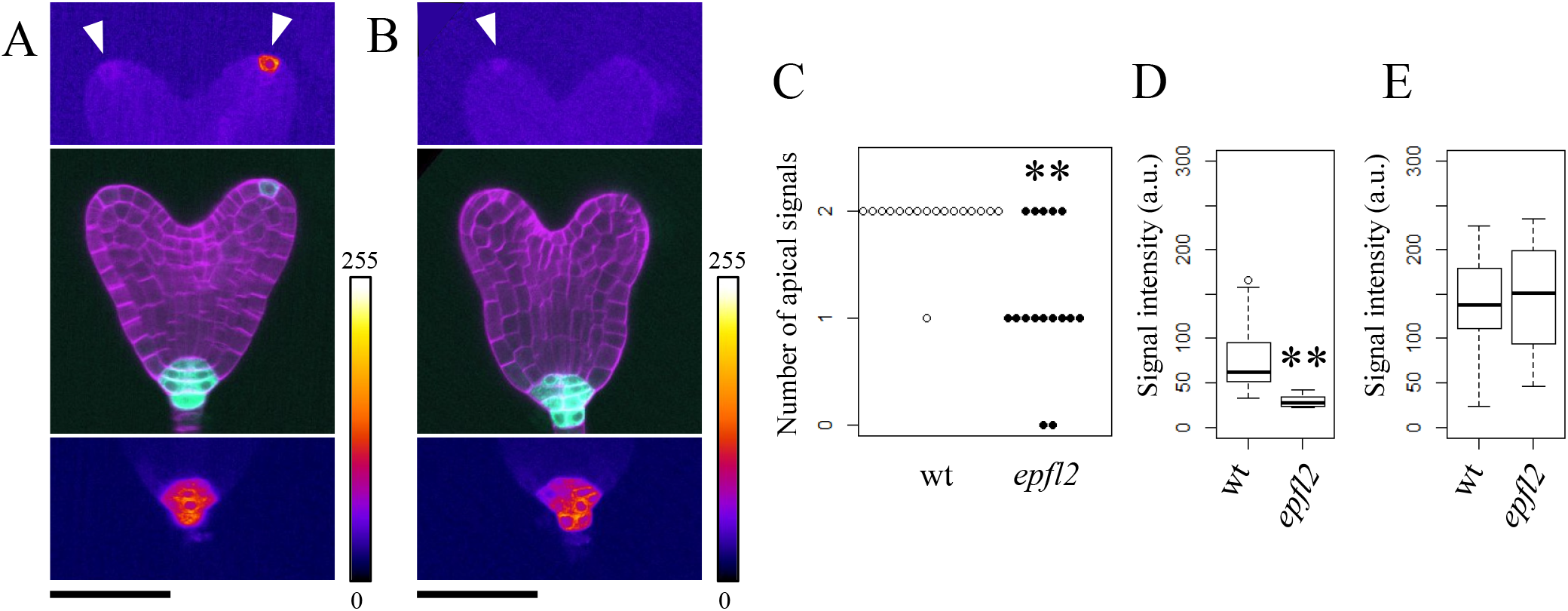
Expression of *DR5rev::GFP*. (A and B) Patterns of *DR5rev::GFP* in the wild type (A) and *epfl2* (B) backgrounds. The whole views of embryos are shown in the middle panels with cell wall staining (magenta) and GFP signals (cyan), with top and bottom panels showing color-coded signal intensities of GFP at the cotyledon (top) and root (bottom) tips of the same embryos. The arrowheads indicate the recognizable GFP signals at the cotyledon tip. (C) Distribution of embryos with different numbers of recognizable apical GFP signals. The open and closed circles represent the individual embryos of wild type and *epfl2*, respectively. (D and E) GFP signal intensity of each genotype in the cotyledon (D) and root (E) tips. The double asterisks indicate the significant differences between the mutant and Col (p < 0.01, Brunner–Munzel test for C and Welch’s t-test for D and E). Scale bars = 50 μm. The sample sizes for each measurement are described in Supplementary File 1.

## Discussion

Our analysis demonstrates that *EPFL2* is required to promote cotyledon growth during embryogenesis. The phenotype of the mutant can be clearly observed from the heart stage, an early stage in cotyledon formation. This result is consistent with the early onset of *EPFL2* expression at the globular stage, indicating that the gene is required for cotyledon growth from its initiation. Our results are also consistent with the non-cell autonomous action of the *EPFL2* gene in cotyledon development, indicating that the cotyledon boundary domain provides a growth-promoting signal by producing the secreted peptide.

The best candidate receptors for EPFL2 are the ERECTA family proteins ER, ERL1, and ERL2, which constitute a subgroup of leucine-rich repeat receptor-like kinases (LRR-RLKs; reviewed by Shpak [2013]). All these proteins have been shown to bind to EPFL2 both *in vivo* and *in vitro*, and genetic studies support that EPFL2 is a ligand for the ER family proteins in leaf tooth growth and ovule patterning (Kawamoto et al. 2020; Tameshige et al. 2016). In addition, the genes for these receptors are all expressed in a broad region that includes the cotyledon primordia in the embryo and are redundantly required for cotyledon growth (Chen and Shpak 2014). To demonstrate that EPFL2 is a ligand for the ER family proteins in cotyledon development, it would be important to test whether the growth-promoting activity of the receptors requires the binding of the secreted peptide. In this regard, it would be also important to test whether the observed difference in the severity of the phenotype between the L*er* and Col alleles of *epfl2* involves the lack of a fully functional *ER* allele in the L*er* background (Torii et al.1996).

An interesting property of *epfl2* mutants is that the reduction in cotyledon size is only apparent in the embryo, and their phenotype recovers by early seedling stage. These results indicate that the *EPFL2*-dependent growth signal is only essential in the early stages of cotyledon development, and the loss of its activity is compensated by other redundant factors. A candidate for such factor is *EPFL1*, an *EPFL* family member that is also expressed in the boundary domain of the cotyledon primordia (Kosentka et al. 2019). In postembryonic development, *EPFL1* is expressed along the boundary between the shoot meristem and leaf primordia and acts redundantly with *EPFL2* and other *EPFL* family members to regulate the shoot meristem size and leaf initiation (Kosentka et al. 2019). Thus, it is likely that the boundary-dependent growth promotion of the cotyledons is supported by multiple redundant factors.

The formation of auxin response peaks is known to be a common key factor in promoting the growth of various organs and tissues (Benková et al. 2003; Bilsborough et al. 2011; Galbiati et al. 2013; Heisler et al. 2005), and the observed reduction in the *DR5* activity at the cotyledon tips of *epfl2* mutants is consistent with this view. However, whether *EPFL2-*dependent signals directly activate the auxin response is still unclear and requires further investigation. It is also important to note that the effect of *EPFL2* on auxin response peaks in the cotyledon tips is significantly different from other developmental contexts. In leaf margin morphogenesis, for example, *EPFL2* rather affects the auxin response negatively and restricts the domain of *DR5* expression to the narrow region within the leaf tooth. In turn, the auxin peak in the tooth represses the *EPFL2* expression, and this mutual repression ensures the spatially complementary patterns of the two factors (Tameshige et al. 2016). A similar negative effect of *EPFL2* on auxin response has also been reported for the vegetative shoot apex (Kosentka et al. 2019). The opposite effects of *EPFL2* on auxin response (positive effect in cotyledons vs negative effect in leaf margins and shoot apices) but the same developmental output (primordium growth promotion) is seemingly paradoxical. However, this can be explained by assuming that the primordium growth is driven by the differential distribution of auxin response rather than its absolute strength–a view that has been shown in leaf margin development (Bilsborough et al. 2011). A comparative analysis of how *EPFL2*-dependent signal regulates the *DR5* expression in different developmental contexts described above will shed light on the variations in the mechanism to establish auxin response peaks by the same signaling peptide.

## Abbreviations

*EPFL*: *EPIDERMAL PATTERNING FACTOR-LIKE*
*A. thaliana*: *Arabidopsis thaliana*
L*er*: Landsberg *erecta*
Col: Columbia
*GUS*: *β- glucuronidase*
CRISPR: clustered regularly interspaced short palindromic repeat
Cas9: CRISPR associated protein 9
GFP: Green Fluorescent Protein
ER: ERECTA
ERL: ERECTA-LIKE
a. u.: arbitrary unit

## Acknowledgements

We thank Maki Niidome, Mie Matsubara, Shoko Nagame, and Kazuko Onga for technical assistance. This work was supported by MEXT KAKENHI (Grant No. 17H06476 to KUT, 24114009, 18H04842, 20H04889 to MA); JSPS KAKENHI (Grant No. JP21H02503 to NU, 16K07401 to MA); WPI-ITbM operational funds to NU and KUT; IROAST operational funds to TH and MA; Takeda Science Foundation to MA.

## Notes

### Competing Interest Statement

The authors have declared no competing interest.

## References

Aida M, Ishida T and Tasaka M (1999) Shoot apical meristem and cotyledon formation during Arabidopsis embryogenesis: interaction among the CUP-SHAPED COTYLEDON and SHOOT MERISTEMLESS genes. Development 126: 1563–1570

Aida M and Tasaka M (2006) Genetic control of shoot organ boundaries. Curr Opin Plant Biol 9: 72–77

Aida M, Tsubakimoto Y, Shimizu S, Ogisu H, Kamiya M, Iwamoto R, Takeda S, Karim MR, Mizutani M, Lenhard M et al. (2020) Establishment of the embryonic shoot meristem involves activation of two classes of genes with opposing functions for meristem activities. Int J Mol Sci 21: 5864

Benková E, Michniewicz M, Sauer M, Teichmann T, Seifertova D, Jürgens G and Friml J (2003) Local, efflux-dependent auxin gradients as a common module for plant organ formation. Cell 115: 591–602

Bilsborough GD, Runions A, Barkoulas M, Jenkins HW, Hasson A, Galinha C, Laufs P, Hay A, Prusinkiewicz P and Tsiantis M (2011) Model for the regulation of Arabidopsis thaliana leaf margin development. Proc Natl Acad Sci U S A 108: 3424–3429

Chen MK and Shpak ED (2014) ERECTA family genes regulate development of cotyledons during embryogenesis. FEBS Lett 588: 3912–3917

Friml J, Vieten A, Sauer M, Weijers D, Schwarz H, Hamann T, Offringa R and Jürgens G (2003) Efflux-dependent auxin gradients establish the apical-basal axis of Arabidopsis. Nature 426: 147–153

Galbiati F, Sinha Roy D, Simonini S, Cucinotta M, Ceccato L, Cuesta C, Simaskova M, Benkova E, Kamiuchi Y, Aida M et al. (2013) An integrative model of the control of ovule primordia formation. Plant J 76: 446–455

Heisler MG, Ohno C, Das P, Sieber P, Reddy GV, Long JA and Meyerowitz EM (2005) Patterns of auxin transport and gene expression during primordium development revealed by live imaging of the Arabidopsis inflorescence meristem. Curr Biol 15: 1899–1911

Hepworth SR and Pautot VA (2015) Beyond the divide: boundaries for patterning and stem cell regulation in plants. Front Plant Sci 6: 1052

Imoto A, Yamada M, Sakamoto T, Okuyama A, Ishida T, Sawa S and Aida M (2021) A ClearSee-based clearing protocol for 3D visualization of Arabidopsis thaliana embryos. Plants 10: 190

Jürgens G and Mayer U (1994) Arabidopsis. In: (Bard, J. ed) Embryos: Color Atlas of Development Wolfe, London pp. 7–21.

Kawamoto N, Del Carpio DP, Hofmann A, Mizuta Y, Kurihara D, Higashiyama T, Uchida N, Torii KU, Colombo L, Groth G et al. (2020) A peptide pair coordinates regular ovule initiation patterns with seed fumber and fruit size. Curr Biol 30: 4352-4361.e4354

Kosentka PZ, Overholt A, Maradiaga R, Mitoubsi O and Shpak ED (2019) EPFL signals in the boundary region of the SAM restrict its size and promote leaf initiation. Plant Physiol 179: 265–279

Palovaara J, de Zeeuw T and Weijers D (2016) Tissue and organ initiation in the plant embryo: a first time for everything. Annu Rev Cell Dev Biol 32: 47–75

Schindelin J, Arganda-Carreras I, Frise E, Kaynig V, Longair M, Pietzsch T, Preibisch S, Rueden C, Saalfeld S, Schmid B et al. (2012) Fiji: an open-source platform for biological-image analysis. Nat Methods 9: 676–682

Shpak ED (2013) Diverse roles of ERECTA family genes in plant development. J Integr Plant Biol 55: 1238–1250

Takeda S, Hanano K, Kariya A, Shimizu S, Zhao L, Matsui M, Tasaka M and Aida M (2011) CUP-SHAPED COTYLEDON1 transcription factor activates the expression of LSH4 and LSH3, two members of the ALOG gene family, in shoot organ boundary cells. Plant J 66: 1066–1077

Tameshige T, Ikematsu S, Torii KU and Uchida N (2017) Stem development through vascular tissues: EPFL-ERECTA family signaling that bounces in and out of phloem. J Exp Bot 68: 45–53

Tameshige T, Okamoto S, Lee JS, Aida M, Tasaka M, Torii KU and Uchida N (2016) A secreted peptide and its receptors shape the auxin response pattern and leaf margin morphogenesis. Curr Biol 26: 2478–2485

Torii KU (2012) Mix-and-match: ligand-receptor pairs in stomatal development and beyond. Trends Plant Sci 17: 711–719

Torii KU, Mitsukawa N, Oosumi T, Matsuura Y, Yokoyama R, Whittier RF and Komeda Y (1996) The Arabidopsis ERECTA gene encodes a putative receptor protein kinase with extracellular leucine-rich repeats. Plant Cell 8: 735–746

Žádníková P and Simon R (2014) How boundaries control plant development. Curr Opin Plant Biol 17: 116–125

