## Supplementary File 1 for "The Boundary-Expressed *EPIDERMAL PATTERNING FACTOR-LIKE2* Gene Encoding a Signaling Peptide Promotes Cotyledon Growth during *Arabidopsis thaliana* Embryogenesis"

**Supplementary File 1. Sample sizes of each mesurements.**

**Figure 2F**

| L*er* | *epfl2-1* |
|---|---|
| 115 | 172 |

**Figure 2G**

| Aixs Height | L*er* | *epfl2-1* |
|---|---|---|
| 51–100 | 75 | 76 |
| 101–150 | 34 | 76 |
| 151–200 | 6 | 20 |

**Figure 2H**

| Col | *epfl2-2* | *epfl2-3* |
|---|---|---|
| 143 | 218 | 233 |

**Figure 2I**

| Aixs Height | Col | *epfl2-2* | *epfl2-3* |
|---|---|---|---|
| 51–100 | 82 | 139 | 124 |
| 101–150 | 45 | 49 | 60 |
| 151–200 | 16 | 30 | 49 |

**Figure 3C**

| Proximo-Distal | Col | *epfl2-2* | *epfl2-3* |
|---|---|---|---|
| Length | 36 | 33 | 37 |
| Cell number | 28 | 34 | 25 |

**Figure 3D**

| Lateral | Col | *epfl2-2* | *epfl2-3* |
|---|---|---|---|
| Length | 79 | 63 | 75 |
| Cell number | 28 | 34 | 25 |

**Figure 4C**

| wild type | *epfl2* |
|---|---|
| 17 | 16 |

**Figure 4D**

| wild type | *epfl2* |
|---|---|
| 17 | 14 |

**Figure 4E**

| wild type | *epfl2* |
|---|---|
| 17 | 16 |
